# Symbiosis of trees and mycorrhiza facilitates displacement of competitors for trees and fungi alike

**DOI:** 10.1101/2020.10.28.358499

**Authors:** Frank Berninger

## Abstract

Ectomycorrhizae are widespread symbionts of higher plants. However, their benefits for plant productivity and growth have not been well demonstrated since many studies do not suggest any improvement of plant growth or of plant nutrition for mycorrhizal plants. We use mechanistic modelling based on the population dynamics of decomposers to simulate the coexistence of mycorrhizal and non-mycorrhizal plants as well as the development of the soil decomposer community. The model assumed a fixed stoichiometry of each decomposer functional type. Decomposer growth depended on its carbon and nitrogen uptake. For mycorrhiza a part of the carbon is modelled to be supplied from the plant while a fixed proportion of the mycorrhizal nitrogen uptake is translocated to the plant. Carbon nitrogen ratios of decomposers were adjusted mineralization of nitrogen or overflow respiration of carbon. The results suggest that mycorrhizal plants do often outcompete non-mycorrhizal plants at no or little improvement of plant productivity.

The main mechanism of mycorrhizal dominance is a reduction of the soil inorganic nitrogen pool and a rerouting of the nitrogen uptake of plants to the transfer nitrogen transfer from mycorrhizae to plants. On the other hand carbon subsidies from the trees allow to expand the niche of mycorrhizal fungi and to outcompete saprohytic fungi under a wide range of physiological and environmental parameters. This leads to dominance of mycorrhizal plants under a broad range of conditions and parameters including low transfer rates of nitrogen from the mycorrhiza to the plant, and low allocation of the plants to mycorrhiza.

**Significance statement:** The paper uses mechanistic, population based modelling to explain the dominance of ectomycorrhizal plants in northern ecosystems while there is limited evidence that these increase plant productivity. We demonstrate that rerouting of the nitrogen cycle towards provision of organic nitrogen to the plant allows mycorrhizal plants to outcompete non-mycorrhizal competitors. Simultaneously, mycorrhizae benefit from carbon subsidies in their competition with saprophytic fungi and bacteria.

## Introduction

Soil ecosystems are locations where plant and diverse decomposer communities are intertwined. Litter from plants is fed to mycorrhiza, saprophytic fungi or soil bacteria which decompose it and make nutrients bound in it available to plants (1). Research shows a network of complex interactions, where both plants and decomposers interact through competition and mutualistic interactions. Mycorrhiza link fungi and plant through a symbiotic relationship, which exchanges nutrients, carbon and water between plants and fungi. Apart from mycorrhizae, other decomposer groups exists as bacteria and saprophytic fungi. These different decomposers have different ecological strategies with bacteria more representing rapidily reproducing ***r***-strategies while fungi are more *K* strategists which expand less aggressively but dominate in stable environments (2). Since decomposers form populations of individuals, we can assume that these populations interact via competition and can be modeled using Lotka-Volterra decomposition like models (3). Furthermore, fungi and bacteria have different stoichiometries with bacteria containing on average three times more nitrogen than fungi. The stoichiometry of decomposers is usually assumed to be constant (4, 5).

Traditional biogeochemical decomposition models like RothC (6) or Century (7) do not include microbes explicitly but organic matter is mineralised using substrate and environment-dependent rates. Newer models that are based on microbial populations and their activity are evolving (3 8, 9). However, only a few of these include mycorrhizae explicitly (8). The more mechanistic of these new biogeochemical models are based on enzyme activity, and decomposer stoichiometry (10, 3). Waring et al. (9) showed that the approach could be extended to simulate the coexistence of functional groups (as fungi and bacteria) during litter decomposition.

Over the past years the utility of mycorrhiza for nutrient uptake has been questioned. Hasselquist et al, (11) as well as Näsholm et al. (12) claim that mycorrhiza contribute little or not at all to the nitrogen nutrition of trees while a recent review shows a positive effect with a large variation of growth in response to mycorrhizal inocculation (13). A possible explanation for the widespread substantial investment of plants into “potentially useless mycorrhiza” comes from research on mutualistic niches. Peay (14) investigated mycorrhizal mutualism through a framework of niche theory and claimed that the mutualistic niche might be larger than the niche of a species in isolation. He, furthermore, stressed that mycorrhizae transform their environment and might extend the niche of their host species. Transformations of the environment by plants or mycorrhiza are mechanisms how mutualistic species help to gain competitive advantages. For example, Corrales et al. (15) suggested that mycorrhiza induced changes in the soil nitrogen cycling might lead to monodominance of the ectomycorrhizal tree species *Oreomunnea mexicana* in humid tropical forests. Also, the competitive displacement was not based on an improvement of the performance of *Oreomunnea* but rather through a reduction of available nutrients and other soil conditions that decrease the performance of the competitors more than the performance of *Oreomunnea*.

Another relevant link between plants, microbes and soil organic matter is an enhanced decomposition of soil organic matter due to carbon inputs from plants to mycorrhiza; a process often called priming. Note that the original definition of priming did not refer to the direct transfer of carbon to mycorrhiza but to rhizodeposition of carbohydrates (16). Allocation of carbon to mycorrhiza could extend the niche of mycorrhizal fungi because it provides energy to acquire nitrogen from soil compounds that might be energetically unfavourable for saprotrophic fungi (17). Priming also increases overall decomposition rates. Lindén et al. (18) and Pumpanen et al. (19) stressed in a mesocosm experiment that the role of plants in the priming process is important since the response of carbon decomposition on sugar additions increased dramatically if plants were present.

In this modelling study, we explore two interrelated questions: (i) How do the different functional types of decomposers: mycorrhizal fungi, saprophytic fungi and bacteria, coexist under varying environmental conditions and (ii) what are the implications of the composition of the decomposer community for nutrient cycling and ultimately plant productivity. Here, we apply an enzyme activity based simple ecosystem model based on the model of Schimel & Weintraub (3) and Waring et al. (9) to simulate ecosystem production, the dynamic of soil organic matter as well as the dynamics of three different functional groups of decomposers.. The model allows for competition between mycorrhizal and non-mycorrhizal plants. Our central hypothesis is that mycorrhizal plants outperform non-mycorrhizal plants due to changes in the carbon and nitrogen supply to the decomposers, leading to lower mineralisation rates of nitrogen. Furthermore, we hypothesize that the turnover of soil organic matter will be faster with mycorrhiza due to priming compared to non-mycorrhizal plants. Also, we hypothesize that low litter carbon/nitrogen ratios in soil organic matter favour non-mycorrhizal decomposer groups, in particular, bacteria (9). High allocation of carbon from trees to mycorrhiza will favour mycorrhizal biomass while high allocation of nitrogen from mycorrhiza to the trees will favour non-mycorrhizal decomposer groups.

### Theoretical modeling framework

We modified the decomposition model (3) inspired by a modified version of Waring et al. (9) who pioneered the inclusion of different functional decomposer groups in Schimel type models. However, neither the Schimel nor the Waring models included a mass balance for the soil organic matter or mycorrhiza. The model has three different functional groups of microorganisms: ectomycorrhizae, saprophytic fungi and bacteria. These differ in their stoichiometry with fungi having a higher carbon to nitrogen ratio than bacteria. Microbial biomass growth is modeled using mass balances of micro-organisms and different microbial functional groups compete for dissolved organic carbon and nitrogen in the soil solution and all microbes as well as plants take up inorganic nitrogen from the soil solution. Mycorrhizae obtain carbon subsidies of the plant and pay a “nitrogen tax” to the plant. Microbes mineralize surpluses in nitrogen and immobilize inorganic nitrogen if there is a surplus of carbon. Surplus carbon is lost in overflow respiration. Soil organic matter is added by litter from plants as well as by dying microbes while it is lost by decomposition. In addition, there are two different plant types mycorrhizal and non-mycorrhizal plants. A simplified flow chart of the model is in figure 1. The model is meant to be conceptual and does not include all plant production, carbon and nitrogen cycling processes.

**Figure 1.**
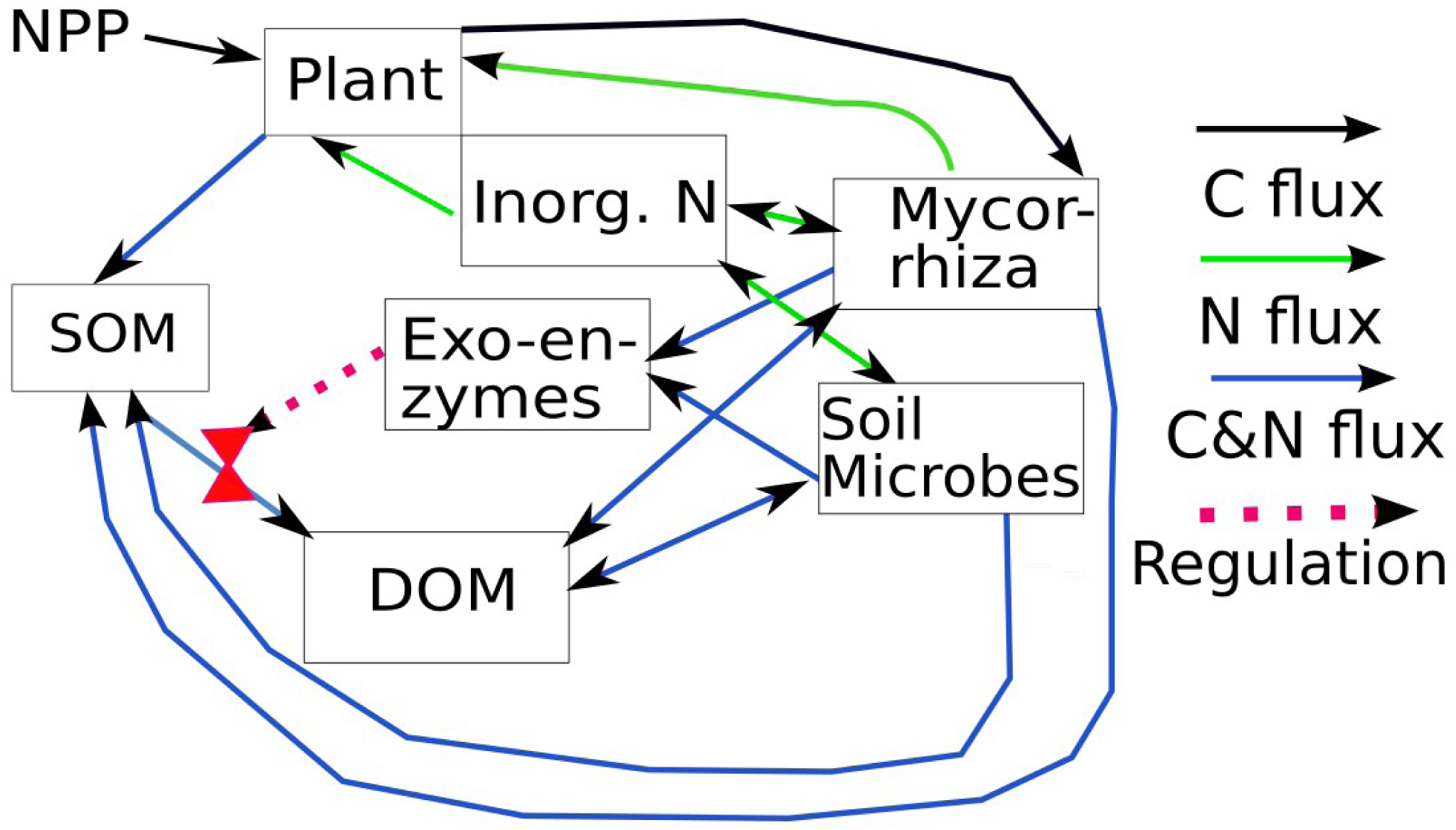
Flow diagram of the model. Saprophytic fungi and bacteria are aggregated under soil microbes to keep the graph readable. Soil and dissolved organic nitrogen and carbon are also combined to keep the figure readable. SOM means soil organic matter, DOM means dissolved organic matter, Nitrogen flows are shown in green, Carbon flows in black and combined carbon and nitrogen flows in blue. The red line indicates that exoenzymes regulate the decomposition of soil organic matter. For simplicity we added only one functional group of soil microbes besides mycorrhiza while the actual model has soil bacteria and saprophytic fungi as functional groups.

## Results

### Comparison with ecosystem development in a post fire chronosequence

We compared the model to published data from a fire chronosequence from a subarctic pine forests in Finland (20, 21, 22, 23) which spanned 160 years of forest development and was sampled at four different ages. The modelling study shows a close correspondence between modeled and measured soil organic carbon. In addition, the development of microbial biomass was similar to observed data (Fig. 1). We found also a decrease in the relative abundance of saprophytic fungi, as has been found by the measurements in the chronosequence (21). There were discrepancies in the C/N ratio of soil organic matter. The observed C/N ratio increased from 23 after fire to 40 about 150 years after the fire while the simulated C/N ratio stabilized at lower values (Fig. 2). The model predicted well a trend for changes in the activities of enzymes involved in nutrient and carbon cycling. Priming accounted for about 12% of the soil organic matter decomposition.

**Figure 2.**
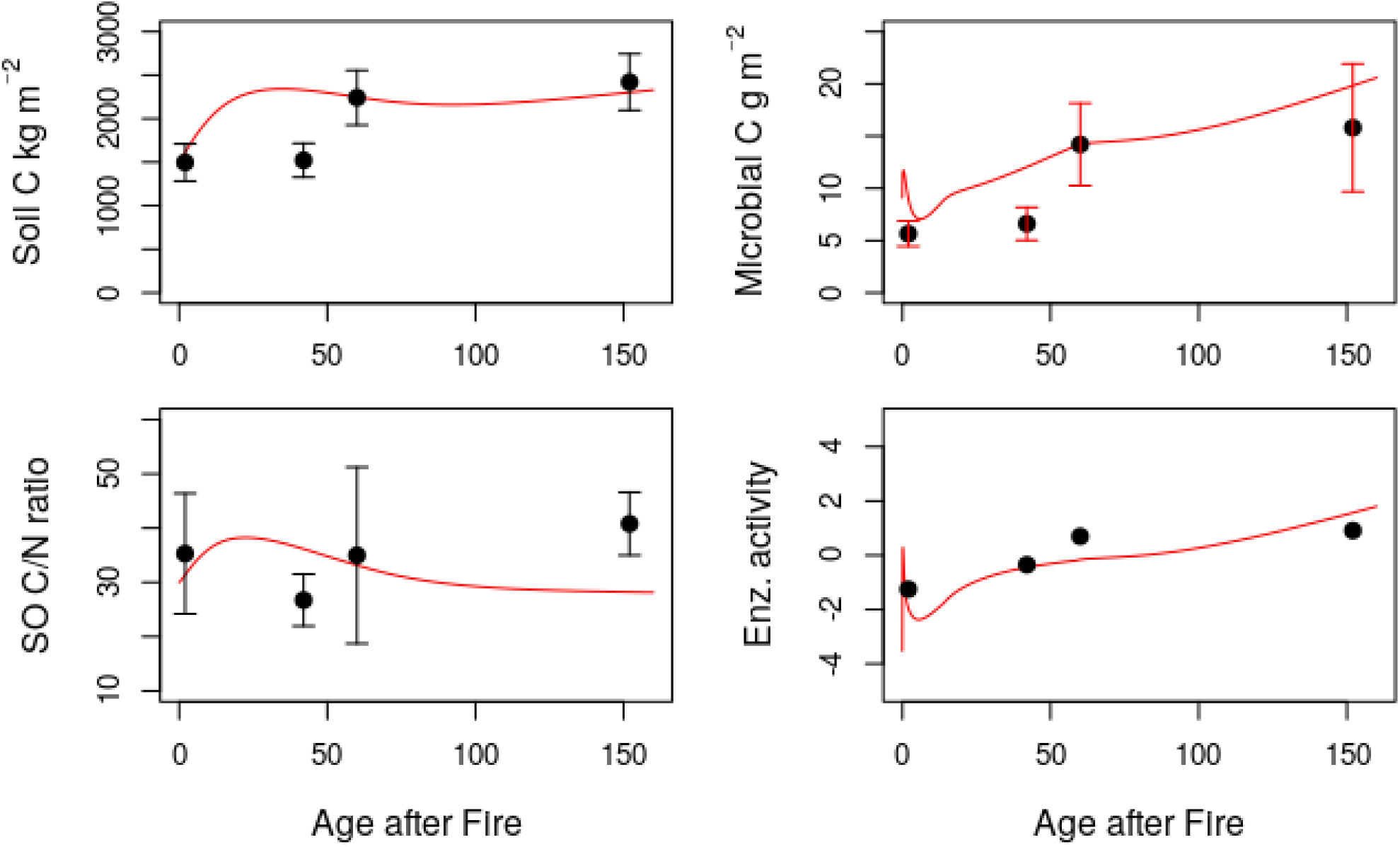
Comparison of the time course of model for the development of ecosystem in Värriö after a non-stand replacing fire. Red lines are modelled values while black dots are measured values. The error bars indicate the standard deviation of the measured values (a) measured and modelled soil organic carbon (g C m^-2^). (b) microbial biomass (g Biomass m^-2^) including bacteria and fungi. (c) Soil organic C/N ratio (g /g). (d) Standardized activity of exoenzymes involved in biogeochemical cycles (unitless).

### Comparison of ecosystems with and without mycorrhizal plants

In a second run we compared the competition of mycorrhizal and non-mycorrhizal plants in a separate run. There were differences in initialisation and parameterisation of this run to the model runs in Fig 2 that are explained in the method section. Fig 3 indicates that the presence of mycorrhiza did not increase the significantly plant biomass or soil organic carbon. However, mycorrhizal plants dominated the ecosystem towards the end of the ecosystem development. Microbial biomass was higher in the presence of mycorrhiza, while nitrogen mineralization rates, as well as the concentration of inorganic nitrogen, were higher in the absence of mycorrhiza. The nitrogen uptake for mycorrhizal plants was initially dominated by uptake of inorganic nitrogen, but the uptake shifted towards organic nitrogen that was translocated from the mycorrhiza.

**Figure 3.**
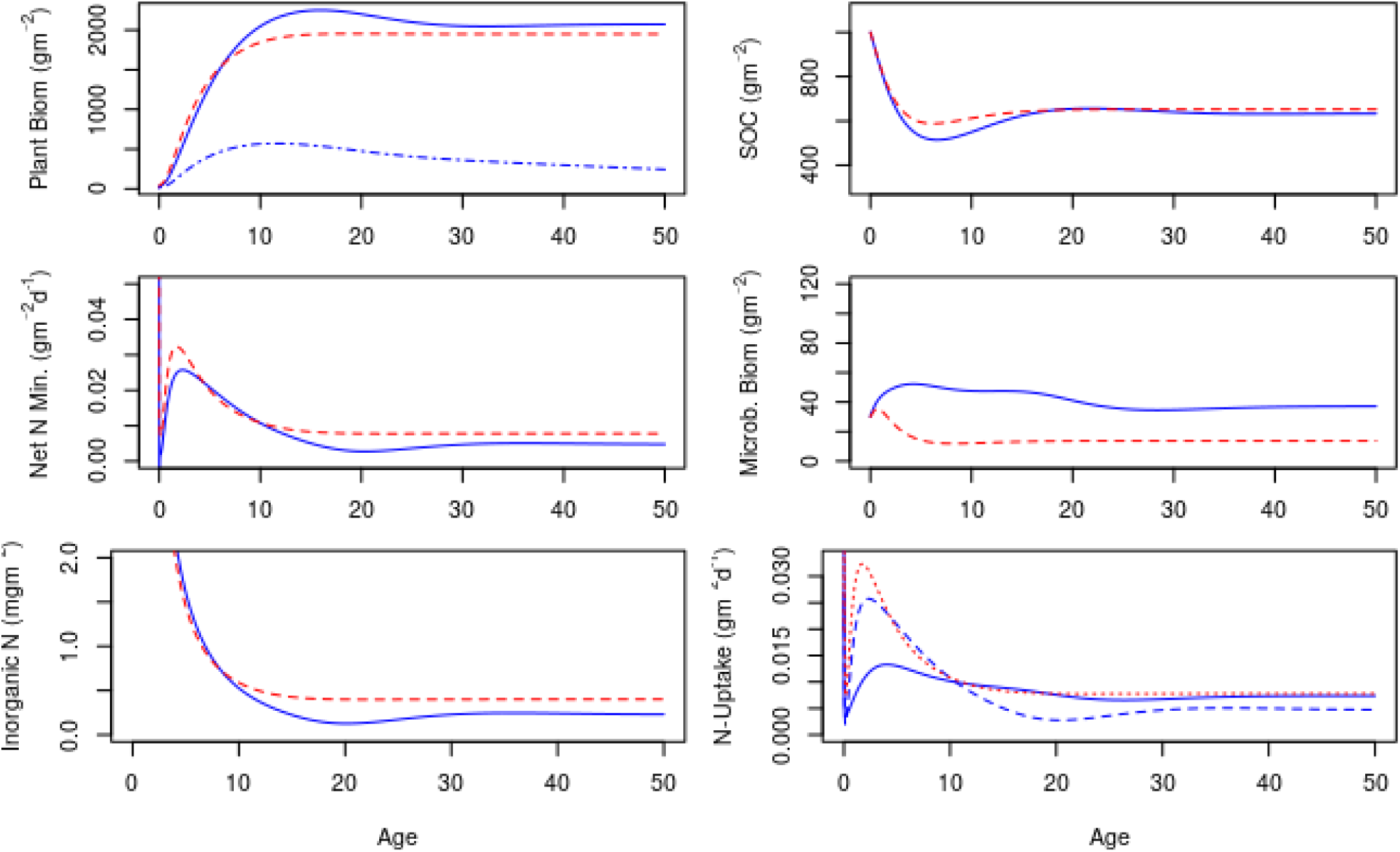
Simulation competition between mycorrhizal and non-mycorrhizal plants. Blue lines depect a system with both mycorrhizal and non-mycorrhizal plants. Red lines depict a system with only non-mycorrhizal plants. (a) Plant biomass (g m^-2^) against time. Red lines denote the biomass for a hypothetical ecosystem with only non-mycorrhizal plant. Blue whole line is the total biomass of plants in an ecosystem that contains both mycorrhizal and non-mycorrhizal plants. The doted blue line is the biomass of non-mycorrhizal plants in the mixed ecosystem. (b) Soil organic carbon (g m^-2^)for ecosystems with mycorrhizal and mixed (mycorrhizal and non-mycorrhizal plants) (c) Concentration of inorganic N in simulated soils (d) N uptake (g of N /day). The whole blue line denotes the total uptake of organic and inorganic N by the plants. The broken blue line the uptake of inorgnaic N only. Red lines denote the uptake of inorganic N in an ecosystem without mycorrhizal plants. The uptake on organic N for non-mycorrhizal plants was assumed to be 0.

### Effects of mycorrhizal N allocation

The second simulation experiment (Fig. 4) explored the effect of changes in the proportion of mycorrhizal N uptake that is allocated to the plant. The values in Fig. 4. presented results are values at steady state after 50 years of simulation. Initially increases in the nitrogen transfer from the mycorrhiza to the plant increase the mycorrhizal biomass, while bacterial biomass and the amount of inorganic nitrogen decrease (Fig. 4a). Also, plant biomass increases (Fig. 4b). At about 20% transfer of the N uptake from mycorrhiza to the plant mycorrhizal and plant biomasses peak while inorganic N stocks in the soil reach a low but stable level. Higher levels of transfer of N from the mycorrhiza to the plant decrease mycorrhizal biomass and the biomass of saprophytic fungi increases while bacterial biomass decreases. Around N transfer rates of about 40% of the N uptake, the biomass of saprophytic fungi decreases as does plant biomass and bacterial biomass increases (not shown). This goes along with a reduction in the carbon nitrogen ratio in the soil (not shown).

**Figure 4.**
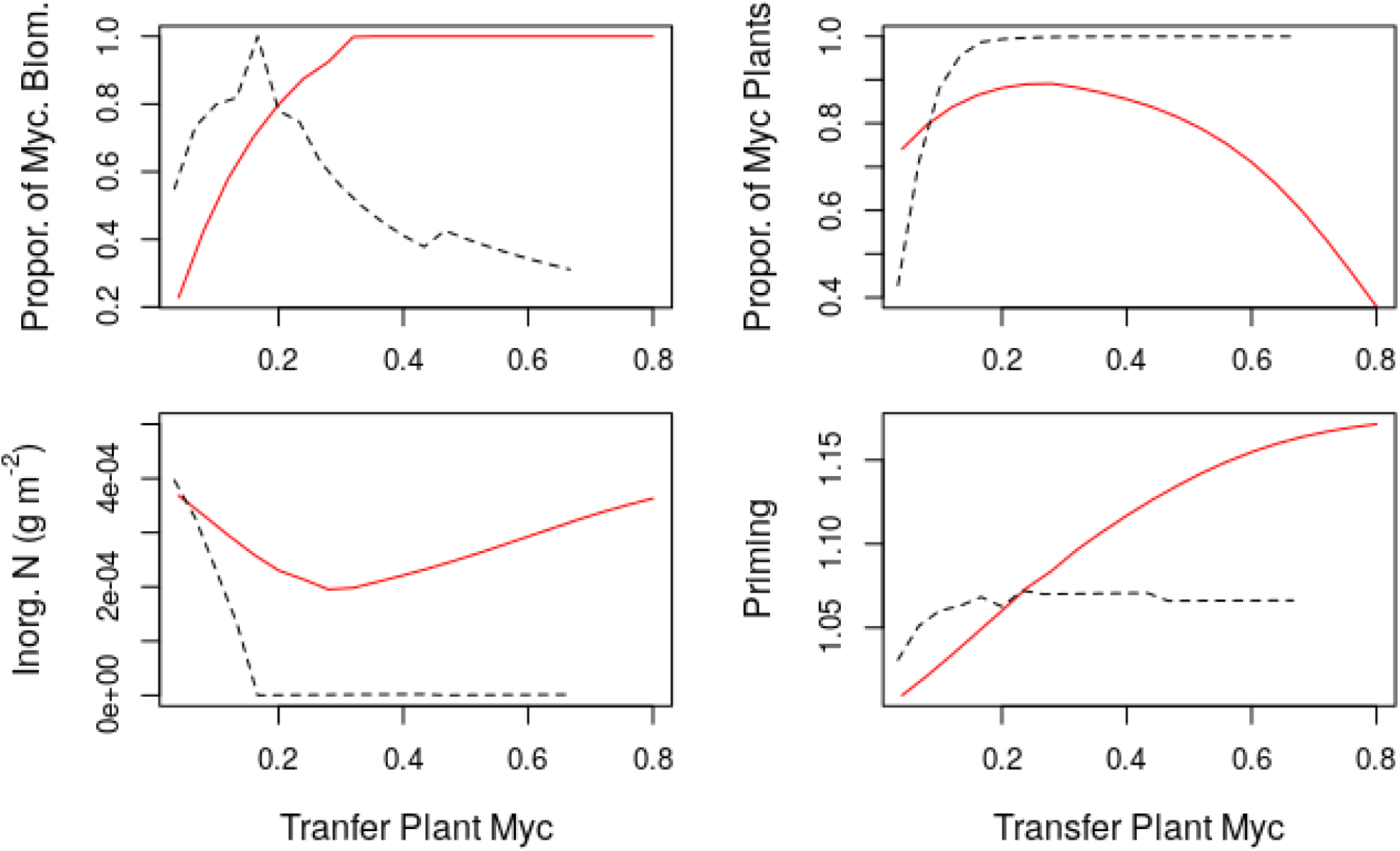
Simulated effects of allocation of plant productivity (red lines) to mycorrhiza and of transfer of N from the mycorrhiza to the plant (black lines). The units of the x-axis are the proportion of net primary productivity that is transferred between mycorrhizal plants and mycorrhiza and the proportion of N uptake from mycorrhiza that is transferred between mycorrhizal plants and mycorrhiza. (a) proportion of mycorrhizal fungi of microbial biomass (b) Proportion of mycorrhizal plants of the whole plant biomass (c) Inorganic nitrogen concentration g m^-2^ (d) Priming, i.e. decomposition caused by allocation of carbon to mycorrhiza. Values or 1 inidicate no priming.

### Effects of plant carbon allocation to the mycorrhiza

As seen in fig. 4 increases in the proportion of Net Primary Productivity (NPP) allocated to mycorrhiza increased the microbial biomass in the soil, usually increasing the mycorrhiza at the detriment of the biomass of bacteria (saprophytic fungi were not abundant in the base run). Above an allocation of about 30% of the NPP to the mycorrhiza plant biomass and soil Carbon/Nitrogen ratio decreased while soil organic matter always decreased with increasing allocation to the mycorrhiza. Also dissolved organic matter (both nitrogen and carbon (not shown) decreased. The proportion of non-mycorrhizal plants decreased with increasing allocation to mycorrhiza, and the biomass of non-mycorrhizal plants did not increase at high allocation to the symbionts, although plant biomass started to decrease. Soil inorganic nitrogen decreased with increasing allocation to mycorrhiza and reached very low values around 30% of NPP-allocated to the mycorrhiza.

### Effects of litter C: N ratio

As seen in Fig. 5, changes in the litter C:N ratio increase the mycorrhizal biomass and decrease the bacterial biomass up to a C:N ratio of about 50, where a stable plateau is reached. They go along with an increase in the soil C:N ratio from low values to values of around 20 while the proportion of non-mycorrhizal plants decreases. Inorganic N concentrations first increase at low litter C:N ratios but then decrease until C:N ratios reach values of about 100. Around litter C:N ratios at 100, bacterial biomass declines while mycorrhizal biomass increases initially, but then decreases while the biomass of saprophytic fungi increases.

**Figure 5.**
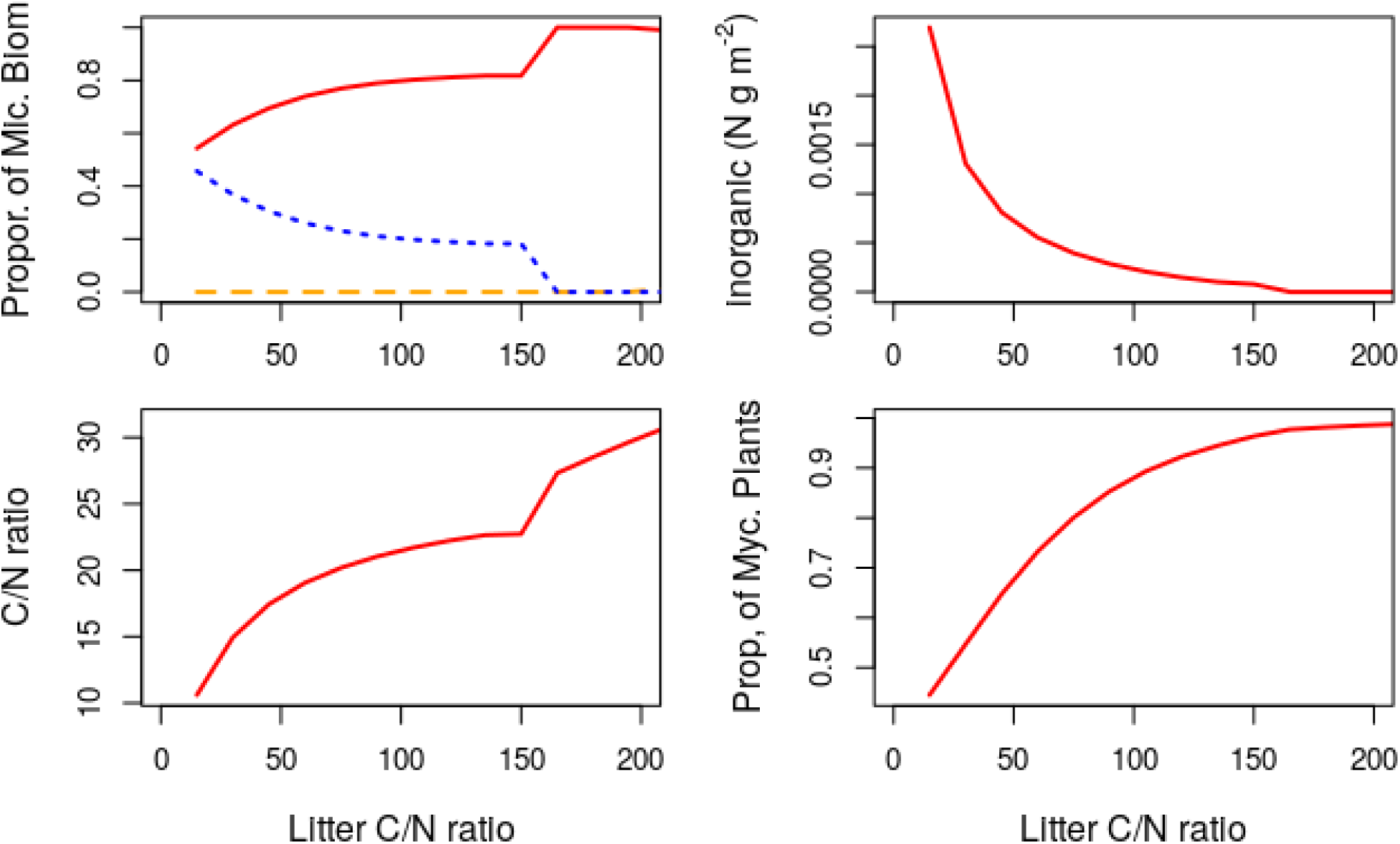
Effects of simulated changes in the plant C/N ration on soil processes. (a) different microbial biomass functional groups (red mycorrhiza), yellow saprophytic fungi and blue soil bacteria. Units are g C m^-2^. (b) inorganic N stocks (g m^-2^) (c) Soil C/N ratio and (d) Proportion of mycorrhizal plants of the biomass.

## Discussion

Our model simulations demonstrate that through modification of the environment, mycorrhizal plants dominate ecosystems while mycorrhizae do not increase ecosystem productivity. We resolve the apparent paradox that mycorrhizae do not facilitate the acquisition of nitrogen (Näsholm et al. 2013) while they have ubiquitous presence of them in northern forest ecosystems. The evolutionary advantage of mycorrhizae are caused by rerouting parts of the nitrogen cycle and by gearing the nitrogen uptake of trees toward organic nitrogen. Organic nitrogen uptake by mycorrhizae provides a competitive advantage to trees because mycorrhizae furnish more efficient access to the organic nitrogen pool that would be otherwise mineralized. However, carbon subsidies from trees to mycorrhiza keep mycorrhizal fungi nitrogen-limited and reduce nitrogen mineralisation rates. This is because carbon transfer from trees increases the need of micro-organisms to acquire nitrogen to meet their stoichionmetry. If carbon supply is low N from mycorrhiza is mineralised.. Therefore, mycorrhizae do not increase ecosystem productivity, although they provide a competitive advantage to mycorrhizal plants. Also, results of our model agree broadly with observed trends over ecological gradients. In the discussion, we describe the main results and compare a few essential emergent results of the model to the literature.

Our model was designed as theoretical model, aiming to understand fundamental processes and not to make quantitative predictions. Nevertheless, our simulations replicate with reasonable skills the development of key ecosystem features in a comparison of the Värriö fire chronosequence. The increases in soil organic matter (20) was similar than in our study. The proportion of mycorrhizal fungi was lower at the beginning of the simulations but increases through time similarly to the observations of (21). About half of the nitrogen uptake of the plants originated from the mycorrhizae which is within the values estimated from Hobbie et al. (24).

We observed discrepancies between measured and simulated Soil carbon nitrogen ratios which tended to continue increasing with time in Värriö while they saturate in the model (20). The soil N stocks decreased during the simulations because an increasing part of the N is stored in the vegetation. This agrees with the observations of Palviainen et al. (23). We think that the differences in modeled and measured carbon/nitrogen ratio in the model can be attributed to a lack of different soil organic matter fractions in the model (like in 6, 7). We decided not to implement these, as they would have resulted in a more complex model and would have made interpretation of our results less straightforward.

Another simplifying assumption is that we assume that there is no direct uptake of organic N by the plant. While there is evidence that plants can take up organic nitrogen directly the ecological importance of this uptake is not well quantified (25). Zhang et al. (26) suggest that myceliar uptake is important for N in mountain forest but claimed to observe an important contribution of direct root uptake of organic N. However, the assumption of no direct uptake of N is also a component of present modeling studies as Ågren et al. (27).

Another discrepancy with observed ecosystem development is that the model shows strong ecosystem tipping points when environmental thresholds are exceeded. This is typical for population based competition models like Lotka-Volterra models. These sharp edges, turning points etc have been described for several population models including stoichiometry driven models as Andersson et al. (28). In real life environmental variability over time and space will definitely lead to more smooth transitions of ecosystem properties along environmental gradients.

Low inorganic nitrogen concentrations in the presence of mycorrhizal trees were the key factors that contributed to the simulated decline of non-mycorrhizal plants in most of our simulation runs. The dominance of mycorrhizal plants occurred over a broad range of parameter values, showing that mycorrhizae increase the fitness of plants over an extended range of environmental conditions, plant and fungal properties. Our results give process-based mechanistic evidence to the idea of Peay (14) that mycorrhizae provide a niche extension of mycorrhizal plants through modification of the environment. In tropical highly diverse mixed forests areas, it has been reported, that ectomycorrhizal trees often form pure monospecific stands (29).

Comparison of model runs using mycorrhizal, and non-mycorrhizal plants show that the presence of mycorrhiza does often not increase the production of plants and does not contribute to a better N nutrition of plants if all plants would be non-mycorrhizal. There has been an ongoing discussion if mycorrhiza does improve the nutrition and growth of plants with many publications demonstrating that mycorrhiza does not increase growth or improve tree N-nutrition. Experimental evidence for this is given, e.g. from Hasselquist et al. (11) who observed that reduction of plant allocation to mycorrhiza resulted in no reduction of N uptake by the plant. Ågren et al. (27) analysed using a similar approach the effect of mycorrhiza on plant growth. They predict that under certain conditions, mycorrhizae might increase plant growth. Their model, differed from our work in scope and assumed a fixed inorganic N concentration and ignored competition between different decomposer groups while it goes beyond this paper in the analysis of mycorrhizal strategies.

Our model attributes low net mineralisation rates of nitrogen to ectomycorrhiza. While the gross mineralisation rates of simulated ecosystems with and without mycorrhizal plants are similar, in mycorrhizal stands immobilisation rates of inorganic N are higher because mycorrhizae are nitrogen limited. The results agree with observed low net nitrogen mineralisation rates in ectomycorrhizal dominated boreal forests. Also, ectomycorrhizal stands show low net mineralization rates and monodominance of ectomycorrhizal species in other ecosystems, as moist evergreen forests. Our results also agree with Blasko et al. (30), who observed that the net mineralization of nitrogen decreased sharply with age after a primary succession chronosequence gets colonized by ectomycorrhizal trees. Also, as in Blasko low net mineralization rates coincide with high microbial biomasses as in our simulations. Studies from agricultural ecosystems also indicate that application of fungicides often increases the net mineralisation of nitrogen (e.g. 31, 32) which is another indication that mycorrhizae cause a decline in net mineralisation of nitrogen. This competition between decomposers and plants has been analysed in detail by Ågren et al. (27), who indicate that mycorrhiza may have a more substantial benefit at high inorganic N concentrations. However, these concentrations were fixed in Ågren’s model while they were a model outcome in our data.

Mycorrhizae were able to dominate the soil microbial community because of the resource (carbon) transfers from trees. These carbon subsidies as well as nitrogen transfers from the fungi to the plant kept mycorrhiza N limited and reduced N mineralization. Mycorrhiza used mostly organic nitrogen pools of which some was transferred to the plants. In the model the switch from the use of organic to inorganic nitrogen in the simulations allows mycorrhizal plants to grab resources and outcompete non-mycorrhizal plants. For example, in our simulations, half of the plant N was taken up via the mycorrhiza at the end of the simulation time (Fig 3). The dominance of organic N as a nitrogen source for the trees agrees with literature (33, 34).

When we modified the proportion of mycorrhizal nitrogen uptake transferred from the fungus to the plant, we observe a rapid decline of the inorganic N concentration because mycorrhizae become increasingly N limited. At the same time, the proportion of mycorrhizal plants increases rapidly. The dominance of mycorrhizal trees is assured at relatively low values of the transfer of N to the trees (around 20%). Also, plant biomass peaks at transfers of N of around 20% of the mycorrhizal N uptake. The reason for the peak in the plant biomass seems that at high values of N transfer, mycorrhizae cannot compete with bacteria for soil resources. Similar results are shown by Ågren et al. (27) who demonstrated that high transfer rates of N from the symbionts to the plant reduce mycorrhizal biomass and, in his study, often ultimately plant growth. The results show that the observed low transfer rates of N from mycorrhiza to trees may have an ecological significance since higher rates of N transfer may not allow mycorrhiza to compete successfully with other soil microbes. Surprisingly, at intermediate levels of N transfer to the tree, there seems to be a distinct niche supporting a higher abundance of soil bacteria. However, the increase in plant biomass in response to increases in N transfer to the plant was relatively modest about 10 % of the biomass. Low allocation of N from mycorrhiza to plant increased the biomass of non-mycorrhizal plants.

Higher allocation of carbon from trees increases initially plant productivity but leads at higher allocation rates to decreases in plant productivity. However, these increases do not seem to open a niche for non-mycorrhizal plants since inorganic N concentrations remain very low. The reason is that N-limited mycorrhizae immobilize inorganic nitrogen and reduce the rates of mineralization of N. Increasing allocation of carbon to mycorrhizae increases first the soil C/N ratio sharply but then leads to a decline. Allocation to mycorrhiza at low and medium levels may lead to small increases in plant biomass while the main effect seems to be on the competitive displacement of non-mycorrhizal plants, saprophytic fungi and soil bacteria. At high proportions of net primary production (NPP) of allocated to mycorrhizae plant biomass declines clearly. We acknowledge that this decline occurs at levels of allocation to mycorrhizae which are higher than values reported in the literature that are usually below 20 % of NPP (34).

Low C/N ratios of litter allow for mineralization of nitrogen and higher concentrations of inorganic N but around a litter C/N ratio of about 50 the levels of inorganic nitrogen drop as has been observed in most boreal forests (which litter C/N ratios are around 50 and above). This coincides with lower C/N ratios in dissolved organic matter as well as with a drop in the bacterial biomass. The model results agree well with observations from nature where ectomycorrhiza dominate in the boreal coniferous forest with low litter C:N ratios but other microbial associations are common in more fertile ecosystems with lower litter C:N ratios. Furthermore, increases in C:N ratios reduce the mineralization of N and make the non-mycorrhizal trees less competitive as proposed by Corrales et al. (15) and other authors (35). Hodge et al. (36) mentioned a threshold of a C:N ratio of 30 where fungi start to immobilize N rather than mineralizing it. It has also been argued that plants on nutrient-poor soils allocate a higher proportion of their NPP to mycorrhiza while the translocation of N from the mycorrhiza to the tree might be lower (17).

Throughout the simulations, the model predicted a modest increase in organic matter decomposition due to allocation of plant NPP to mycorrhiza. The effect was typically less around 10% of the total decomposition of soil organic matter. The priming increased with increasing allocation to mycorrhiza. We point out that the definition of priming in this publications is more conservative to experimental definitions (where sugar is added to the soil) since experimental definitions include direct supply of C to non-symbiotic organisms, changes in the microbial community structure and increase of microbial populations due to priming (16).

We aimed to present here a simple, parsimonious model that allows the analysis of the grand picture of the interaction of ectomycorrhizae and plants. Central simplifications that might affect the model outcome are the lack of variations in organic matter recalcitrance which is driving most standard decomposition models as RothC. Another simplifying assumption is that different decomposers use a homogenous carbon and nitrogen pool in the same way. Research using stable isotope probing demonstrates that decomposers differ in their nutrient and carbon sources (37) assumptions of a fixed C:N ratios of decomposer groups, simplified assumptions on plant growth and the assumption that non-mycorrhizal plants are completely unable to take up nitrogen.

The present modelling is based on a number of assumptions that are presently under active discussion. Firstly, we assume that the C/N ratios of each decomposer functional group are fixed, while changes in the microbial C/N ratio are caused by changes in the microbial community (4). For example, Deckmyn et al. (38) give large ranges of variations of mycorrhizal C/N ratios. Variation in fungal C/N ratio within the microbial population would probably smooth edges in the figures 3-6. Furthermore, it is possible that microbial stoichiometry reacts to the environment. However, Zhou et al. (5) suggest that the stoichiometry of boreal microbial communities does not respond to changes in elemental ratios of the substrate.

Secondly, we assume that organic N-uptake is limited to soil microbes and that organic N-uptake is not possible for plants. This is partly a modelling assumption to allow for a comparison of an idealized mycorrhizal and an idealized non-mycorrhizal plant. While it is known that plants can take up certain types of inorganic N, the quantitative role of inorganic N that plant take up directly is under discussion (e.g.). We acknowledge that if mycorrhizae do not provide a more efficient way of acquiring organic nitrogen than roots, our model would not work and not address relevant questions.

Our plant production model assumes a constant Nutrient Use Efficiency and a constant C/N ratio of litter throughout ecosystem development. Vitousek (1) realized the relationships between Nitrogen Use Efficiency (NUE) and the C/N ratio (or in his case the N content) of litterfall. He also showed that constant NUE models work reasonably well as estimators of productivity across large spatial scales. We acknowledge that over a succession the nutrient use efficiency of boreal forest will increase with time (41), Since we aim, at this stage of model development, to understand processes across broad spatial and temporal scales we did not attempt to include varying NUE into our model.

Furthermore, our model does not describe soil organic matter quality in terms of different soil organic matter compartments that are too different degrees recalcitrant to decomposition as in most decomposition models. Lindahl & Tunlid (17) outline that carbon and nitrogen in the most recalcitrant forms could be accessible predominantly to mycorrhiza since the decomposition process is exothermic.

## Conclusions

In our theoretical modeling study, we suggest that mycorrhiza might provide competitive advantages to plants by routing plant nutrition from inorganic to organic forms and reducing mineralization of inorganic nitrogen. Mutualistic advantages allow mycorrhizal plants to suppress the growth of non-mycorrhizal competitors and to outcompete non-mycorrhizal fungi but do not go along with significant increases in plant productivity.

## Materials and Methods

A detailed technical description, including equations of the model, as well as the R-code to run the model, are published in appendix 1.

Plant net primary production (NPP) (including growth allocated to mycorrhiza) is assumed to be proportional to the plant nitrogen uptake. The ratio of plant production and nitrogen uptake equals the carbon nitrogen ratio of the litter following the ideas of (1) assuming for simplicity a constant Nitrogen Use Efficiency which equals the carbon per nitrogen ratio of the litter input. We define two different plant functional groups (mycorrhizal and non-mycorrhizal plants). We assume that organic nitrogen is not taken up by plants directly. However, mycorrhizal plants may access nitrogen via transfer from the fungi. Plant senescence was assumed to be a fixed proportion of plant biomass and this biomass returns to the soil as litter. Both mycorrhizal and non-mycorrhizal plants take up soil inorganic nitrogen as a fixed proportion of the inorganic nitrogen stock. Mycorrhizal plants receive, in addition, a fixed proportion of the nitrogen taken up by mycorrhiza and that is used for growth. This includes organic and inorganic nitrogen taken up by the mycorrhizae.

The Schimel and Weintraub model is a model that simulates litter decomposition as a function of enzyme activity and decomposer stochiometry. The model assumes that the stochiometry of decomposers is constant (4). Decomposers produce exo-enzymes that degrade soil organic matter and produce dissolved organic matter. Decomposers take up this dissolved organic matter for growth and respirations. Excess carbon is respired using overflow respiration while excess nitrogen is mineralized. Ammonium and nitrate are not modelled explicitly, and no transformations of inorganic nitrogen as Nitrification are modelled. Microbes may also acquire nitrogen via uptake of inorganic N called here immobilisation.

Our modification of the model was to divide the decomposer pool into saprophytic fungi, mycorrhizal fungi and bacteria. These functional groups have different C/N ratios, as reported by Waring et al. (9)). The C/N ratio for bacteria was set to 5, while the ratio for fungi was set to 15 (9). We differ from the Waring paper by assuming that microbial dissolved organic matter uptake is proportional to the share of each functional group in biomass and the activity of the functional group, while Waring assumed it to be proportional to the share of the enzyme production of each functional group.

In the model, priming is defined as the ratio of the decomposition of soil organic matter that is caused by the transfer of carbon from the plant to the mycorrhiza to the decomposition that would occur without support from the plant (see equation 26 in the supplementary material). This definition of priming is instantaneous, i.e. it excludes changes of microbial biomass or community structure.

### Technical implementation

The model is implemented as a differential equation system under R. The model has 11 state variables and 28 parameters (of which many were set to identical values for the different functional groups). Simulations were done for 5000 days using the “lsoda” solver (40). Integration was done using the deSolve package (41). The solver determines the simulation time step, but the output is set to 1 day.

### Base parameterization

Key parameter values are the C/N ratios of the different functional groups that were set to 15 for fungi (both saprophytes and mycorrhiza) as well as to 5 for bacteria (9), For the base run 20% (34) of the NPP or mycorrhizal plants was allocated to mycorrhiza while 15% of the N uptake from mycorrhiza was transferred to trees. We assumed that bacteria had a 4 times higher organic carbon and nitrogen uptake than fungi (on a mass basis) due to a higher cell surface to cell volume ratio (42). Litter C/N ratio was set to 120 for the base run, while the initial soil C/N ratio of the soil was 20 (23). The turnover of the bacteria, i.e. the proportion of bacteria dying each time step, was 0.035/day while it was 0.009/day for fungi (9). The senescence for plants was set to 0.0003/day (43). We assumed that the carbon uptake of bacteria per unit of mass is 4 times the rate for fungi since they have a higher mass to surface ratio. The parameter values are representative for a northern boreal forest ecosystem dominated by ectomycorrhiza.

### Comparison with real data

We based our simulation on the microbial biomasses carbon stocks as well as initial carbon/nitrogen ratios of a fire chronosequence in Northern Finland (Värriö) where we have done extensive studies of soil processes (20). We set initial biomasses, soil carbon stocks and carbon nitrogen ratios to the observed values. Also total microbial biomasses were set to the values of Köster et al. (20) by scaling measured microbial C concentrations to the whole soils. The run differs from the simulation experiments described below by the addition of nitrogen net deposition based on values reported in Palviainen et al. (23). For the simulation experiments nitrogen deposition was assumed to be zero. Enzyme activities were are the average of the activity of soil enzymes involved in biogeochemical cycles reported by Köster et al. (23). Since direct comparison with simulated soil enzyme concentrations is not possible we compared standardize values. Standardized enzyme activities were calculated by calculating for each enzyme the mean activity over time (for both simulated and measured enzyme activities).

### Simulation experiments

We organized our simulations around five simulation experiments that explore the effects of mycorrhiza on ecosystem processes. Simulations were performed for 6000 days. Note that the growing season at the Värriö site used for much of the parameterization is about 120 days, and our simulations would, therefore, cover around 50 years of ecosystem development

1) We first explored there the effects of mycorrhiza on productivity. We compared the development of the model over time of an with and without mycorrhizal plants. The non-mycorrhizal system was simulated by setting the initial values for mycorrhizal plant biomass and the mycorrhizal biomass to 0. The initial value of the total fungal biomass was kept constant by adding the mycorrhizal biomass to the biomass of saprophytic fungi. Simulations with mycorrhizal plants always included non-mycorrhizal plants at initialization. We investigated the temporal development of microbial biomasses and other simulated variables.
For the simulation experiments, we investigate the behaviour of the model at steady state (after 5000-time steps)
2) The second experiment explores the varying allocation of mycorrhizal obtained N for growth of mycorrhiza and plants. The percentage of N-transfer from mycorrhiza to plants was varied between 5% and 95% while the percentage of plant NPP allocated to mycorrhiza was kept constant (at 20%).
3) The third experiment explored the allocation of plant NPP to mycorrhizae, that was varied between 4% and 80% while keeping all other variables as in the base run.
4) The fourth experiment varied the plant litter C/N ratio, which was varied between 7.5 and 150. It should be acknowledged that ecosystem productivity varies, since plant productivity is modelled as the product of litter C/N ratio and plant nitrogen uptake.

